# A point mutation in the kinase domain of CRK10 leads to xylem vessel collapse and activates defence responses

**DOI:** 10.1101/2021.08.16.456532

**Authors:** Maiara Piovesana, Ana K. M. Wood, Daniel P. Smith, Michael J. Deery, Richard Bayliss, Esther Carrera, Johnathan A. Napier, Smita Kurup, Michaela C. Matthes

## Abstract

Cysteine-rich receptor-like kinases (CRKs) are a large family of plasma membrane-bound receptors ubiquitous in higher plants. They are transcriptionally regulated by a wide variety of environmental cues and stresses, however their precise biological roles remain largely unknown. Here we report a novel mutant isolated for the CYSTEINE-RICH RECEPTOR-LIKE KINASE 10 (CRK10) of *Arabidopsis thaliana* which harbours the substitution of alanine 397 by a threonine in the αC-helix of its kinase domain and which we registered as *crk10-A397T* in the community database. *In situ* phosphorylation assays with the His-tagged wild type (WT) and *crk10-A397T* versions of the CRK10 kinase domain revealed that both alleles are active kinases capable of auto-phosphorylation with the newly introduced threonine acting as an additional phosphorylation site in *crk10-A397T*. Phenotypically the mutant is a dwarf and the analysis of thin cross sections with light and transmission electron microscopy revealed that collapsed xylem vessels in roots and hypocotyls are very likely the cause for this reduction in stature. Transcriptomic analysis of WT and mutant hypocotyls revealed that predominantly biotic and abiotic stress-responsive genes are constitutively up-regulated in the mutant. Root-infection assays with the vascular pathogen *Fusarium oxysporum* demonstrated that the *crk10-A397T* mutant has enhanced resistance to this pathogen compared to WT plants. Taken together our results suggest that *crk10-A397T* is a gain-of-function allele of *CRK10* and open up new avenues for the investigation of this elusive receptor-like kinase family.

## INTRODUCTION

Plant growth and development is modulated by intrinsic growth regulators as well as by beneficial and adverse environmental cues. Factors regulating development and environmental and pathogenic signals are mostly recognised by receptor-like kinases (RLKs), membrane-localized receptors which perceive and transduce these signals inside plant cells to activate programmes directing appropriate responses. Similar to animal receptor tyrosine kinases (RTKs), these receptors consist of an extracellular domain which perceives specific ligands, a single pass transmembrane domain, and a cytoplasmic kinase domain which transduces the signal via phosphorylation of downstream target proteins in the cytoplasm in order to tailor a cellular response (Shiu & Bleecker, 2001; De Smet et al., 2009). Their highly variable extracellular domains are used to classify RLKs into subgroups, the largest of which (∼200 genes in *Arabidopsis thaliana*) being characterised by leucine-rich repeats (LRR-RLKs) (Diévart & Clark, 2004). The well-studied brassinosteroid receptor BRASSINOSTEROID INSENSITIVE 1 (BRI1; AT4G39400) and the microbial pattern recognition receptors (PRR) FLAGELLIN SENSING 2 (FLS2; AT5G46330) and EF-TU RECEPTOR (EFR; AT5G20480) belong to this subgroup (Friedrichsen et al., 2000; Chinchilla et al., 2006; Zipfel et al., 2006).

Among the several subfamilies of RLKs found in plant genomes, the cysteine-rich receptor-like kinases (CRKs) is one of the largest with over 40 members in *Arabidopsis*. The signature motif for CRKs is the presence of, in most cases, two repeats of the DOMAIN OF UNKNOWN FUNCTION 26 (DUF26) in their extracellular domain, which contains three cysteine residues in the conserved configuration C-X8-C-X2-C (Chen, 2001). The functional significance of the DUF26 domain remains to be elucidated but was originally suggested to participate in ROS or redox sensing (Wrzaczek et al., 2010; Bourdais et al., 2015). However, more recent data obtained from the crystallographic analysis of the DUF26-containing ectodomain of the plasmodesmata localised proteins (PLDPs) PLDP5 and PLDP8 point towards the involvement of the cysteine residues in forming disulfide bonds for the structural stabilisation of the protein, rather than redox sensing (Vaattovaara et al., 2019). The same study also revealed that the DUF26 domain shows strong structural similarity to fungal carbohydrate-binding lectins, which suggests that DUF26-containing proteins might constitute a group of carbohydrate-binding proteins in plants (Vaattovaara et al., 2019). That DUF26-containing proteins do interact with carbohydrates has been shown for the secreted antifungal protein from *Gingko biloba*, Ginkbilobin2 (Gnk2), which contains a single DUF26 domain and acts as a mannose binding lectin (Miyakawa et al., 2009; Miyakawa et al., 2014), and for the secreted antifungal proteins AFP1 and AFP2 from maize, which contain tandem DUF26 domains also binding mannose (Ma et al., 2018). Ligands for the CRKs, however, still remain to be discovered.

Despite the prominence of the CRKs among the RLKs, very little is known about their specific functions and the regulation of downstream signalling events. Efforts to assign functions to CRKs involved a comprehensive analysis of a collection of T-DNA knockout lines for 41 CRKs of Arabidopsis, which suggested a role for several members in the fine-tuning of stress adaptation and plant development. Most knockout lines, however, did not display obvious phenotypes, as is expected for a large gene family due to redundancy amongst its members (Bourdais et al., 2015). Studies in Arabidopsis also revealed that several CRKs are transcriptionally regulated by a wide variety of biotic and abiotic factors such as ozone, ultraviolet light (UV), reactive oxygen species (ROS), the hormone salicylic acid (SA) and elicitation with pathogen-derived molecules (Czernic et al., 1999; Du & Chen, 2000; Ohtake et al., 2000; Wrzaczek et al., 2010). Functionally, CRKs belong to the RD subclass of Ser/Thr kinases (Vaattovaara et al., 2019), which typically carry a conserved arginine (R) immediately preceding the invariant aspartate (D) in subdomain VI required for catalytic activity and are in most cases activated through auto-phosphorylation of the activation loop (Nolen et al., 2004). Although the ability to auto-phosphorylate as well as to phosphorylate substrates *in situ* has been demonstrated for CRK2 and CRK7 (Idänheimo et al., 2014; Kimura et al., 2020), for example, detailed studies on the *in vivo* and *in vitro* role of phosphorylation sites are still outstanding for CRKs.

Gain-of-function mutations of a kinase are a valuable tool to study specific effects on cells and organisms, as in these instances the signalling cascade is initiated spontaneously without requiring the presence of an extracellular ligand. Although much is known about the structure–function properties of eukaryotic kinases, predicting which point mutations will lead to a spontaneous activation of the signalling cascade is not straightforward and mutations are generally fortuitously isolated from mutagenic screens. In this study we describe the characterisation of such a gain-of-function mutant, *crk10-A397T,* obtained for the cysteine-rich receptor-like kinase CRK10 (AT4G23180) of *Arabidopsis.* This mutant contains a single amino acid substitution from alanine to threonine in the αC-helix of the kinase domain of the protein, with the newly introduced Thr-397 acting as an additional *in situ* phosphorylation site. Defense responses are constitutively activated in the *crk10-A397T* mutant, as detected by transcriptome profiling, and resistance to a soil-borne vascular pathogen is significantly enhanced. In addition, we could link the dwarf phenotype of the mutant to the severe collapse of the xylem vessel elements which we showed to occur in the root and hypocotyl but, surprisingly, not in the shoot.

## RESULTS

### *crk10-A397T* is a semi-dominant mutant allele of CRK10

The *Arabidopsis thaliana* mutant characterised in this report was isolated in a forward genetic ethyl methanesulfonate (EMS) screen performed for an unrelated study. In brief, six rounds of backcrosses to the wild-type (WT) Col-0 parent were performed in order to clean the genetic background of the mutant before an in-depth characterisation. The homozygous mutant has a strong dwarf phenotype and observation of the segregating F2 progeny of the sixth backcross revealed the semi-dominant nature of the mutation, as WT, intermediate and dwarf phenotypes segregated according to a 1:2:1 ratio with heterozygous plants being clearly discernible (Supplemental Figure S1A; 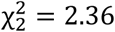, p = 0.308). In order to determine the underlying mutation responsible for the dwarf phenotype, whole genome sequencing was performed on bulk segregants derived from the sixth backcross. This returned a list of 15 candidate genes containing point mutations in coding regions. We noticed that a substitution of G>A in the 4th exon of the *CYSTEINE-RICH RECEPTOR-LIKE KINASE 10* (*CRK10*; AT4G23180) lead to the substitution of alanine 397 by a threonine in the kinase domain of CRK10 (Supplemental Figure S1B). As we considered this receptor-like kinase to be the most likely candidate among the 15 identified genes, we tested whether the dwarf phenotype could be rescued by constitutive expression of the WT cDNA sequence of *CRK10* under the control of the 35S promoter. All T1 transformants showed a WT phenotype (Supplemental Figure S1C), suggesting that we had identified the correct gene. To further confirm that the mutation in *CRK10* causes the dwarf morphology, we recreated the G>A substitution by *in vitro* mutagenesis in the cDNA sequence of *CRK10*, and introduced this ORF into a *crk10* KO background (*crk10-2*, SAIL_427_E09, characterisation of KO lines to follow) under the control of the 1 kb genomic region containing the putative native promoter of *CRK10.* A total of 25% of the recovered transformants were dwarfs, establishing a direct link between the dwarf phenotype and the mutant allele (Supplemental Figure S1D). Subsequently, we will refer to this mutant as *crk10-A397T*.

### The A397T substitution is localised in the αC-helix of the kinase domain of CRK10

According to the subdivision of eukaryotic kinase domains into 12 conserved subdomains (Hanks & Hunter, 1995), the A397T substitution is localised in subdomain III of the kinase domain of CRK10 (Figure 1A), which corresponds to the αC-Helix motif in the three-dimensional structure of the protein. Homology modelling to the active kinase domain of the Arabidopsis BRASSINOSTEROID INSENSITIVE 1 (BRI1) positions Thr-397 at the C-terminal end of the helix, with its side chain likely to be exposed on the surface of the protein (Figure 1B).

**Figure 1.**
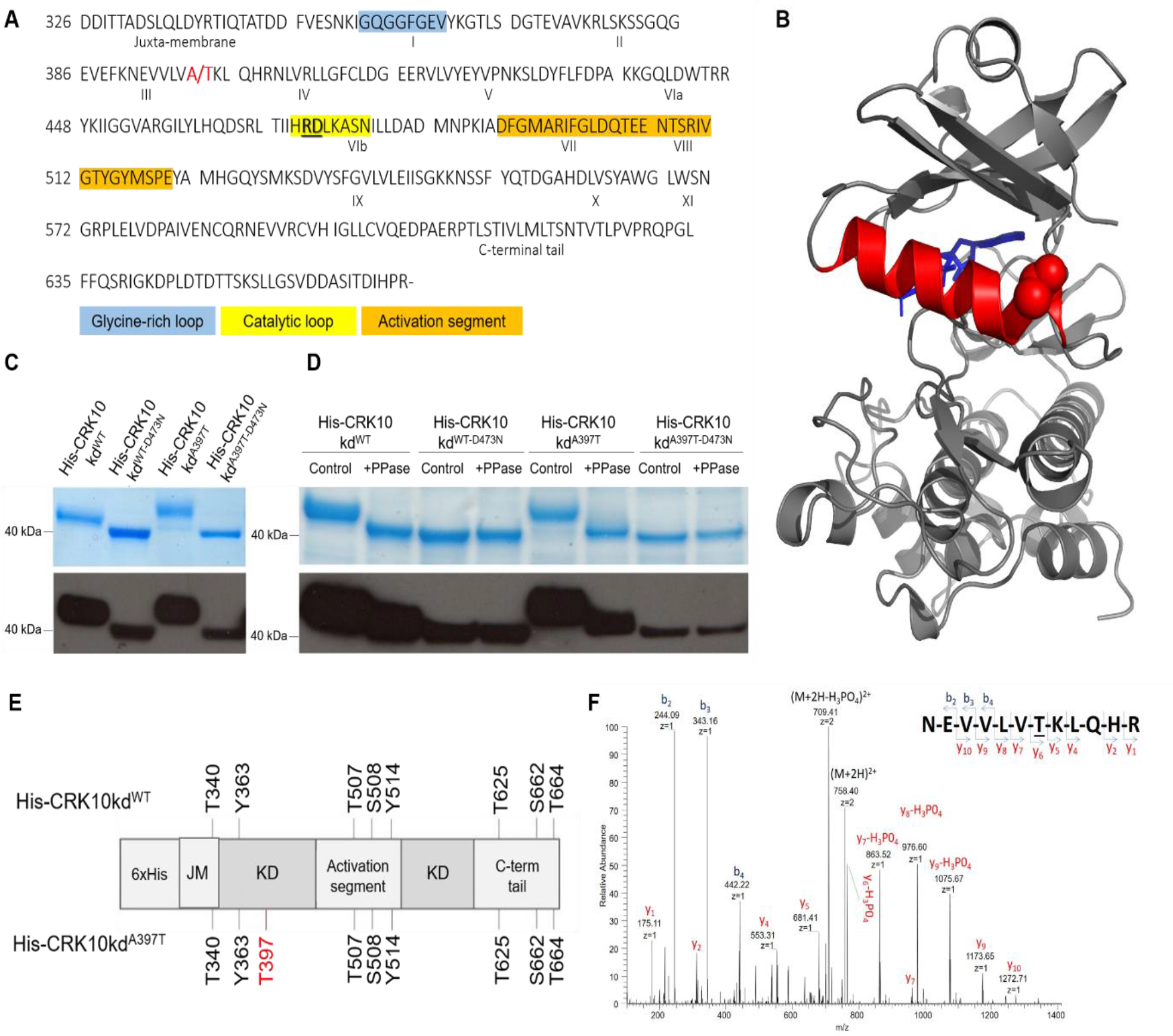
CRK10 and CRK10-A397 are active kinases and the A397T substitution introduces an additional auto-phosphorylation site in the kinase domain of the protein. A, Amino acid sequence of the cytoplasmic kinase domain of CRK10 used for in situ phosphorylation studies (CRK10 amino acid residue numbering shown on the left). Subdomains I-XI are indicated by roman numerals. Ala/Thr 397 site is highlighted in red, and the RD motif is underlined and highlighted in bold. The conserved glycine-rich loop, catalytic loop and activation segment motifs are also shown. B, Structure of the CRK10 kinase domain generated by homology modelling to the active kinase domain of BRI1. A molecule of an ATP analogue, coloured blue, occupies the active site. The αC-helix is coloured red, and the atoms in the side chain of the mutated threonine residue are depicted as red spheres. C, Recombinant His-CRK10kd^WT^ and His-CRK10kd^A397T^ extracts and their respective dead kinase controls (His-CRK10kd^WT^-D473N and His-CRK10kd^A397T-D473N^) resolved by SDS-PAGE; detection by Western blot with HRP-conjugated anti-His antibody is shown. D, SDS-PAGE and Western blot following treatment of purified His-tagged proteins with λ-phosphatase (PPase); control: untreated protein. E, In situ auto-phosphorylation sites in His-CRK10kd^WT^ and His-CRK10kd^A397T^ identified by LC-MS/MS analysis of the recombinant protein kinase domain. Threonine 397 is highlighted in red. His: 6x His-tag; JM: juxta-membrane domain; KD: kinase domain; C-term tail: C-terminal tail. (f) MS/MS spectrum of the doubly charged (m/z 758.4) tryptic phosphopeptide NEVVLVTKLQHR in which the threonine residue is phosphorylated. Neutral losses of phosphoric acid from both the precursor ion and the C-terminal y ions are observed.

### The cytoplasmic kinase domains of CRK10 and CRK10-A397T are active kinases

With CRKs being classified as Ser/Thr kinases, the substitution of Ala-397 by a threonine (A397T) in the CRK10 kinase domain could have introduced a potential additional phosphorylation site. We therefore wanted to determine whether WT and mutant CRK10 are enzymatically active kinases and if differences in their auto-phosphorylation pattern can be detected. We adressed this question by investigating the auto-phosphorylation activity of the cytoplasmic kinase domain of CRK10 *in situ* when expressed in *Escherichia coli* cells (Taylor et al., 2013). We purified the CRK10 WT cytoplasmic kinase domain as an N-terminal 6x His-tag fusion protein (His-CRK10kd^WT^) from *E. coli* cells, as well as its “dead” kinase counterpart which harboured the substitution of the essential aspartic acid 473 with an asparagine residue (His-CRK10kd^WT-D473N^). Following separation of the recombinant proteins by SDS-PAGE and detection by anti-His immunoblotting, the dead kinase version His-CRK10kd^WT-D473N^ migrated at the predicted molecular weight (*M_r_*) of 40 kDa, while the WT kinase version His-CRK10kd^WT^ showed an electrophoretic mobility shift to larger *M_r_*, known to occur for phosphorylated proteins (Wegener & Jones, 1984) (Figure 1C). In order to determine whether the A397T substitution in the CRK10 kinase domain affects its auto-phosphorylation activity, we generated constructs in which Ala-397 was replaced by threonine through *in vitro* mutagenesis (His-CRK10kd^A397T^ and His-CRK10kd^A397T-D473N^). Although the dead kinase version His-CRK10kd^A397T-D473N^ migrated at the same *M_r_* as His-CRK10kd^WT-D473N^ on SDS-PAGE gels, the mobility shift of the His-CRK10kd^A397T^ was increased when compared to the one observed for His-CRK10kd^WT^ (Figure 1C), suggesting additional sites were phosphorylated in His-CRK10kd^A397T^. In order to confirm that the electrophoretic mobility shift was due to the presence of phosphorylated residues, the purified recombinant proteins were treated with λ-phosphatase prior to separation by SDS-PAGE. Irrespective of the treatments, the dead kinase versions CRK10kd^WT-D473N^ and His-CRK10kd^A397T-D473N^ migrated at the predicted *M_r_* as confirmed by SDS-PAGE and anti-His immunoblotting (Figure 1D). However, λ-phosphatase treatment of His-CRK10kd^WT^ and His-CRK10kd^A397T^ resulted in a clearly detectable shift to a lower *M_r,_* consistent with the auto-phosphorylation of recombinant His-CRK10kd^WT^ and His-CRK10kd^A397T^ as being responsible for their electrophoretic mobility shift. Taken together, these results confirm that both His-CRK10kd^WT^ and His-CRK10kd^A397T^ are active kinases capable of auto-phosphorylation. Furthermore, the increased mobility shift of His-CRK10kd^A397T^ compared to His-CRK10kd^WT^ suggested the presence of additional phosphorylation sites in His-CRK10kd^A397T^.

### The kinase domain of CRK10 auto-phosphorylates highly conserved residues in the activation loop and Thr-397 is an additional phosphorylation site in His-CRK10kd^A397T^

We next proceeded to identify which sites in the kinase domain of CRK10 are being phosphorylated by subjecting tryptic peptides of His-CRK10kd^WT^ and His-CRK10kd^A397T^ to analysis by liquid chromatography-tandem mass spectrometry analysis (LC-MS/MS). The Mascot probability-based algorithm was used to confirm the peptides match to the CRK10 kinase domain sequence. Individual MS/MS spectra were inspected for confirmation of phosphorylation sites, which led to the unambiguous identification of Thr-340, Tyr-363, Thr-507, Ser-508, Tyr-514, Thr-625, Ser-662 and Thr-664 as phosphosites in both His-CRK10kd^WT^ and His-CRK10kd^A397T^ proteins (Figure 1E). Interestingly, Thr-507, Ser-508, and Tyr-514 align to conserved phosphorylation sites in the activation loop of several RLKs, known to be essential for the activation of RD kinases (Supplemental Figure S2). Phosphorylated residues were also detected in the juxta-membrane region (Thr-340) as well as in the C-terminal tail of CRK10 (Thr-625, Ser-662, and Thr-664), which are predicted to act as regulatory sites for interaction with binding partners. In addition, the identification of two phosphorylated tyrosine residues (Tyr-363 and Tyr-514) classifies CRK10 as dual specificity kinase and constitutes the first instance in which such activity has been reported for a CRK. Interestingly, Thr-397 itself was identified as a phosphorylation site in the His-CRK10kd^A397T^ kinase domain *in situ* (Figure 1F), however, whether Thr-397 also acts as a phosphorylation site *in vivo* remains to be determined.

### The *crk10-A397T* mutant is a dwarf

WT Col-0 and *crk10*-A397T plants were phenotypically characterised for the duration of one life cycle. Although germination rate and establishment of seedlings is accelerated in the mutant (Supplemental Figure S3), one week after sowing these differences are no longer apparent. No other obvious differences in growth were observed between WT and *crk10*-A397T seedlings until between weeks two and three, when leaf expansion becomes restricted in the *crk10-A397T* mutant as small, dark green leaves are formed, and a reduction of more than 70% in rosette size is observed at four weeks after sowing compared to the WT (Figure 2A-G; Supplemental Figure S4A). Despite the dwarf phenotype displayed during vegetative growth, the onset of flowering occurs simultaneously in *crk10-A397T* mutant and WT plants, although the shoot apical meristem is frequently aborted in the mutant (Supplemental Figure S4B), and the main inflorescence remains stunted (Figure 2H). At later stages, mutant plants develop numerous lateral inflorescences with smaller, stunted siliques filled with viable seeds which are in general larger than those of WT plants (Figure 2-IK).

**Figure 2.**
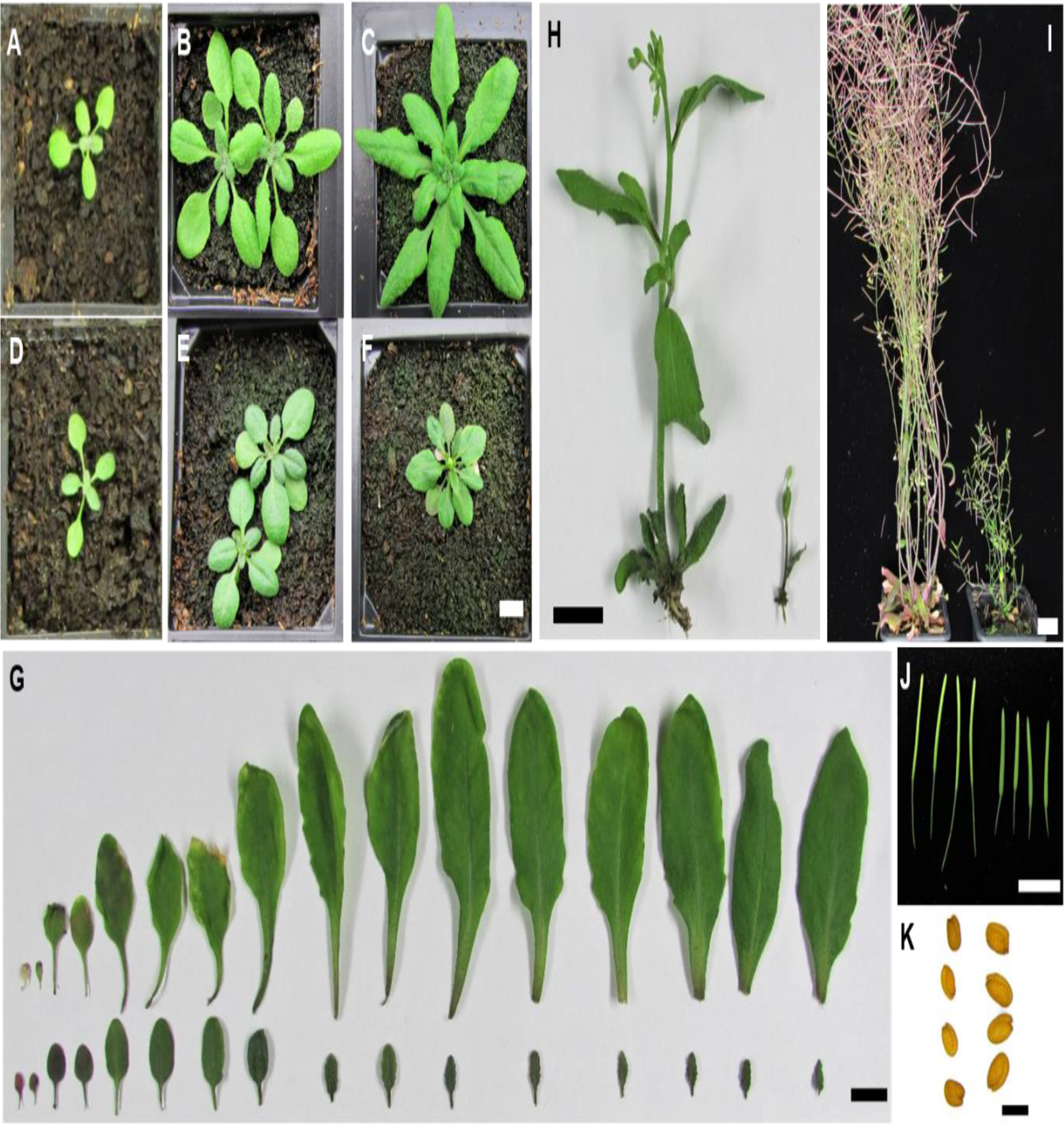
*crk10-A397T* mutant is a dwarf. A-F, Rosette morphology of WT (A, B, C) and *crk10-A397T* (D, E, F) plants at 2 (A, D), 3 (B, E) and 4 (C, F) weeks after sowing. Bar = 1 cm. G, Leaf series of 4-week-old WT (top) and *crk10-A397T* (bottom) plants. Bar = 1 cm. H, Main inflorescence stem of 5-week-old WT (left) and *crk10-A397T* (right) plants. Bar = 1 cm. I, 10-week-old WT (left) and *crk10-A397T* (right) plants. Bar = 2 cm. J, Siliques of WT (left) and *crk10-A397T* (right) plants. Bar = 1 cm. K, Seeds of WT (left) and *crk10-A397T* (right) plants. Bar = 500 µm.

### The *crk10*-A397T mutant has collapsed xylem vessels in roots and hypocotyls

Dwarfism in plants is often caused by defects in the vascular system. To investigate whether the vasculature of the *crk10-A397T* mutant develops normally, we prepared transverse cross sections of resin-embedded hypocotyl, root and stem samples of 5-week-old plants. The sections were stained with potassium permanganate, a lignin-specific dye which allows the observation of lignified xylem vessels and fibres. Imaging of the cross sections revealed that xylem vessels in the root and hypocotyl of the mutant plants are severely collapsed, whereas the vasculature in the stem is not morphologically altered (Figure 3A-F). Interestingly, the collapsed xylem vessels displayed darker brown staining in response to the dye compared to their WT counterparts, which suggests that their secondary cell walls are more heavily lignified (Figure 3G-H). This hypothesis was reinforced by the visualization of auto-fluorescence of lignin of these cells using confocal microscopy, as the xylem vessels in the mutant hypocotyl consistently exhibited a more intense auto-fluorescence signal compared to the WT (Figure 3I-J). Transmission electron microscopy (TEM) was used to investigate potential defects in the ultrastructure of the secondary cell wall of the collapsed xylem vessels in 3-week-old hypocotyls, but no differences were observed in electron density and general appearance of the cell walls compared to that of intact vessels in the WT (Supplemental Figure S5).

**Figure 3.**
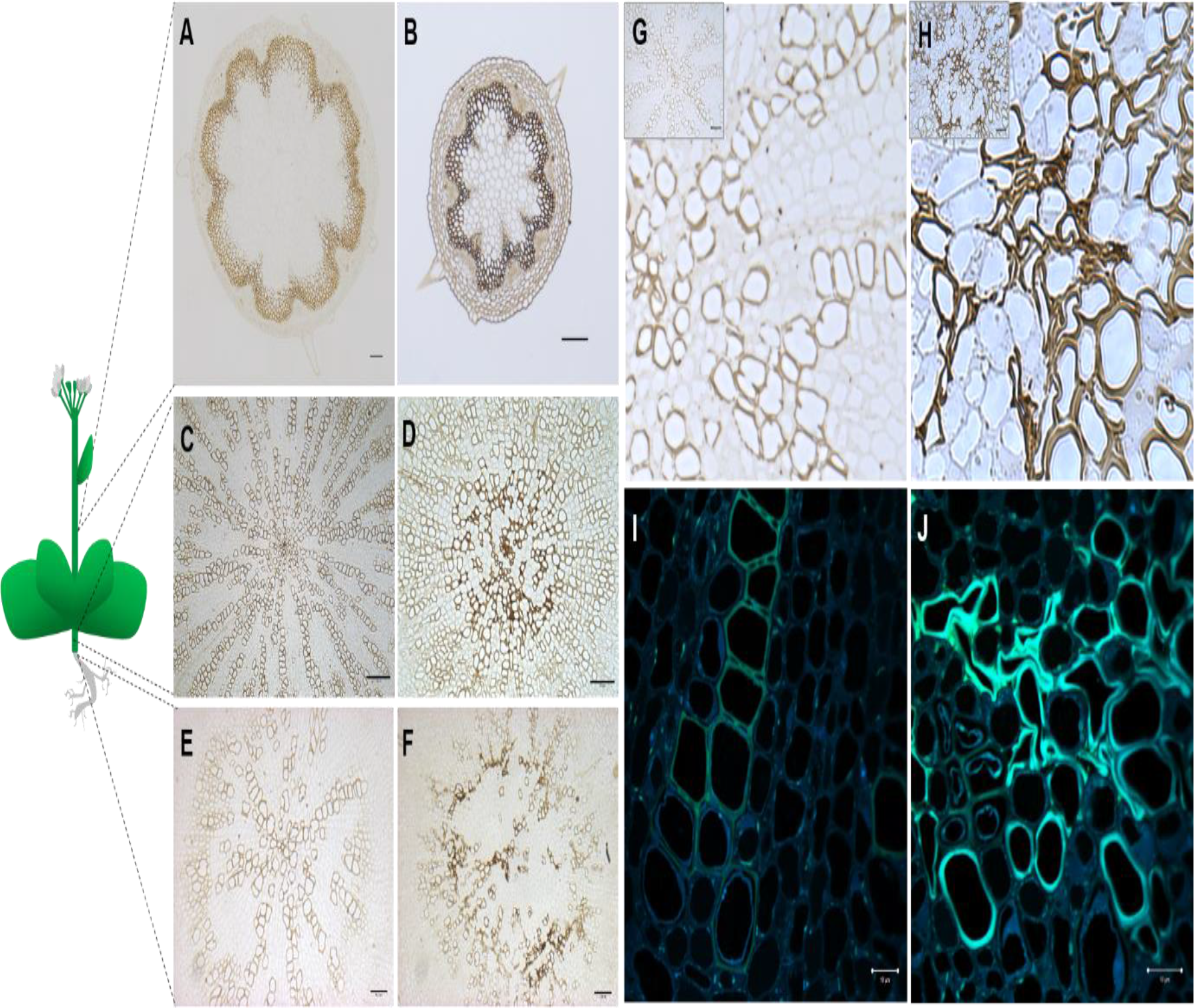
Xylem vessels collapse in the root and hypocotyl of *crk10-A397T* plants, but not in the stem. A-F, Transverse cross sections of the base of stem (A, B), hypocotyl (C, D) and roots (E, F) of 5-week-old WT (A, C, E) and *crk10-A397T* (B, D, F) plants. Stain: potassium permanganate. Bars = (A, B, C, E, F) 100 µm; (D) 50 µm. G-H, Detail of xylem vessels in hypocotyls of 4-week-old WT (G) and *crk10-A397T* (H) mutant plants. Stain: potassium permanganate. Insert in top left corner of image shows original micrographs. Bars = (insert G) 50 µm; (insert H) 25 µm. I-J, Detection of auto-fluorescence of lignin on resin-embedded cross sections of 4-week-old hypocotyls of WT (I) and *crk10-A397T* (J) plants. Bars = 10 µm.

To understand the progression of the phenotype, a developmental time series of hypocotyl cross sections spanning weeks one to five after sowing was analysed. Cross sections of one- and two-week-old hypocotyls revealed the disorganization of xylem vessels in *crk10-A397T* plants at early developmental stages, as they do not proceed to form the typical radial patterning observed in the hypocotyl vasculature of WT plants (Supplemental Figure S6A-D). At three weeks of age, the first deformed and collapsed xylem vessels become apparent in the mutant hypocotyls, a phenotype which is even more severe in 4-week-old plants (Supplemental Figure S6E-H). At 5 weeks of age, following the onset of flowering, cross sections revealed the absence of fully differentiated xylem fibres in the hypocotyl of the *crk10-A397T* plants, in contrast to the WT (Supplemental Figure S6I-L). Differentiation of xylary fibres in *Arabidopsis* hypocotyls is tightly associated with the switch to growth phase II of xylem development, which is triggered by the transition to flowering. We conclude that this switch is delayed in the *crk10-A397T* plants, despite flowering occurring simultaneously to WT plants.

### Loss of function or overexpression of *CRK10* does not cause a phenotype in Arabidopsis

In order to investigate whether increased levels of *CRK10* expression has observable phenotypic effects, we introduced a construct carrying the WT cDNA of *CRK10* under the control of the constitutive 35S promoter in WT *Arabidopsis* plants. Two independent homozygous lines were generated and selected for further analysis (*CRK10* OE-1 and OE-2). Compared to WT, qPCR performed on 4-week-old leaves detected a *CRK10* transcript increase of 15 and 6 times for *CRK10* OE-1 and OE-2, respectively, although growth and development were not altered (Supplemental Figure S7). In order to investigate whether the absence of the *CRK10* transcript affects the phenotype of Arabidopsis plants, two homozygous T-DNA knockout lines for the *CRK10* gene were isolated, *crk10-2* (SAIL_427_E09) and *crk10-4* (SALK_116653). Quantification of *CRK10* transcript levels from leaves of 4-week-old plants by qPCR confirmed that *crk10-2* and *crk10-4* are a knockout and knockdown line of *CRK10*, respectively, however growth and development of both lines were indistinguishable from WT plants (Supplemental Figure S8). Cross sections of hypocotyls of 4-week-old *crk10-2* and *CRK10* OE-1 plants were imaged to rule out the presence of collapsed xylem vessels (Supplemental Figure S9). Lines *CRK10* OE-1 and *crk10-2* were used for the pathogen assays which are described in a subsequent section.

### *CRK10* is expressed in close association with vascular tissues and the protein localises to the plasma membrane

Tissue specific expression of *CRK10* was determined by placing the reporter *β-glucuronidase* under the control of the 1 kb genomic sequence containing the putative promoter of CRK10 (*CRK10*_Pro_:GUS). *GUS* expression was detected in the vasculature of the roots, cotyledons, petioles, leaves, hypocotyls and inflorescence stem (Figure 4A-B; Supplemental Figure S10A-D). In hypocotyls of 2-week-old seedlings, expression was localised to differentiating xylem vessels and parenchyma cells surrounding the vessel elements (Figure 4B). The presence of the *CRK10* transcript in hypocotyls and inflorescence stems of 3- and 6-week-old WT and *crk10-A397T* mutant plants, respectively, was confirmed by qPCR (Supplemental Figure S10E).

**Figure 4.**
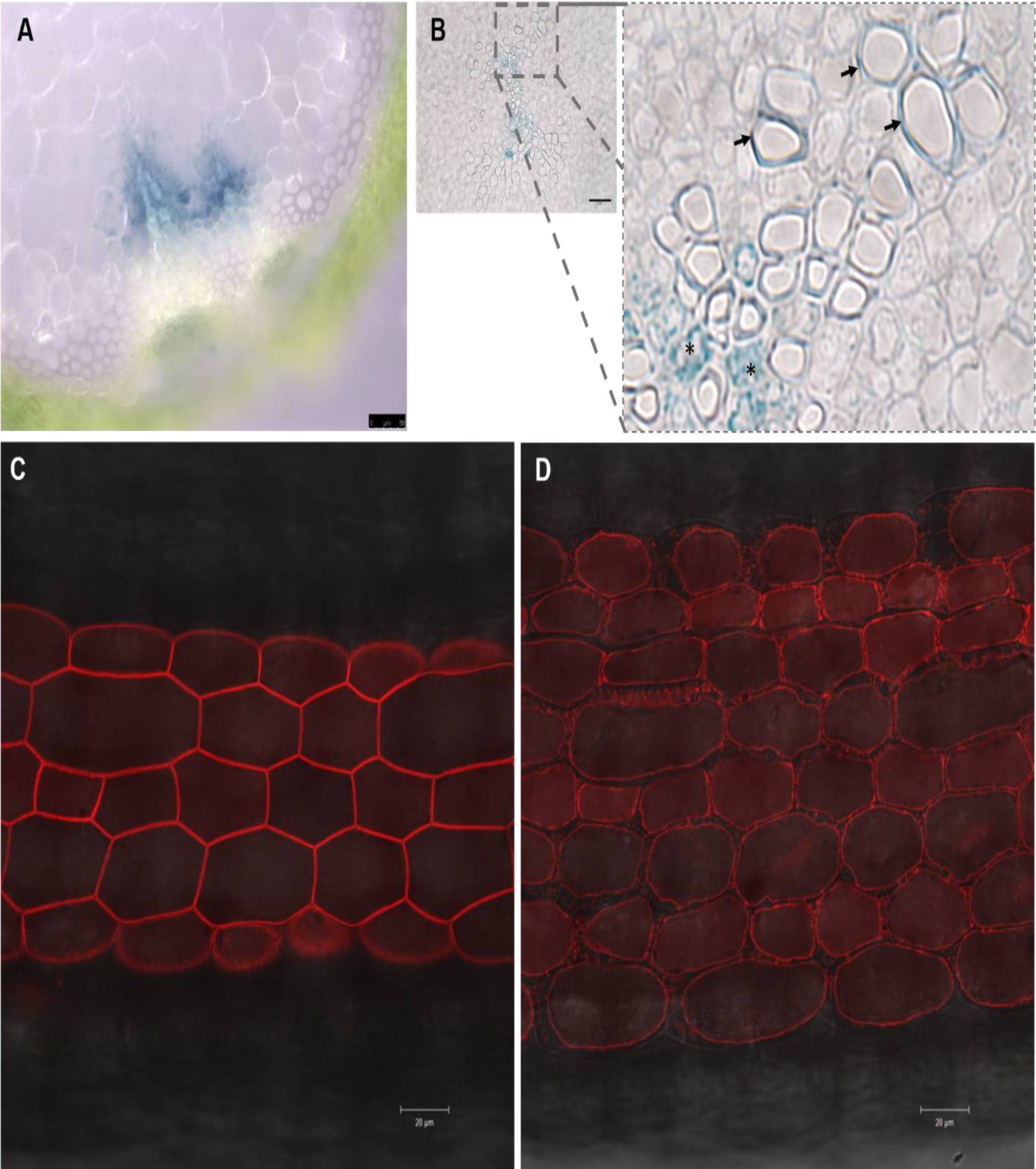
CRK10 is a plasma membrane-localised protein expressed in association with the vasculature in the hypocotyl. A-B, Histochemical staining of reporter lines expressing the *CRK10*_Pro_:*GUS* construct showed expression of the reporter gene in the vasculature of stem and hypocotyls, as shown by free-hand cross section of 8-week-old inflorescence stem (A) and cross section of 2-week-old hypocotyl embedded in resin (B). Bars = (A) 50 µm; (B) 25 µm. C, Detail of cross section in B shows presence of histochemical staining in xylem parenchyma cells (asterisks) and differentiating xylem vessels (arrows) in the hypocotyl. C-D, Hypocotyl of 4-day-old seedling of transgenic Arabidopsis plant expressing 35S:*CRK10-mCherry* before (C) and after (D) plasmolysis. Bars = 20 µm.

Subcellular localisation of CRK10 was determined by analysing lines expressing a construct carrying the C-terminal translational fusion of *CRK10* with the fluorescent protein mCherry under the control of the constitutive 35S promoter (*35S*:*CRK10*-*mCherry*). Both transient expression of the construct in *N. benthamiana* leaves and stable expression in transgenic *Arabidopsis* plants indicated that the fusion protein localises to the plasma membrane (Supplemental Figure S10F; Figure 4C). The presence of Hechtian strands, characteristic of the retracting plasma membrane from the cell wall following plasmolysis (Oparka, 1994), further confirmed this subcellular localisation of the protein (Figure 4D). Therefore, we conclude *CRK10* is expressed in close association with vascular tissues of below and aboveground organs, and the protein localises to the plasma membrane of plant cells.

### Collapsed xylem vessels in the root and hypocotyl are responsible for the dwarf phenotype of the *crk10-A397T* mutant

Although *CRK10* is expressed in tissues associated with vasculature in the stem, hypocotyl and roots, as demonstrated by C*RK10*_Pro_:*GUS* analysis, it is intriguing that in the *crk10-A397T* plants xylem vessel collapse occurs only in roots and hypocotyls. To investigate if the dwarf phenotype is solely due to the defects of the below ground tissues, or whether it is a “whole-plant” response, we performed a micrografting experiment (Turnbull et al., 2002). *In vitro* grown 4-day-old seedlings were used to generate combinations of WT rootstocks and *crk10-A397T* scions (WT/*crk10-A397T*) and vice-versa (*crk10-A397T*/WT), as well as self-grafted plants as controls (WT/WT and *crk10-A397T*/*crk10-A397T*). Phenotypic assessment of successful grafts revealed that a WT scion grafted onto a *crk10-A397T* rootstock develops the characteristic dwarf phenotype of the mutant, whereas a mutant scion develops into a WT-like plant when grafted onto a WT rootstock (Figure 5). Our observations show that the root and hypocotyl system of the *crk10-A397T* plants are responsible for their dwarf phenotype, which is likely due to the presence of collapsed xylem vessels in these tissues.

**Figure 5.**
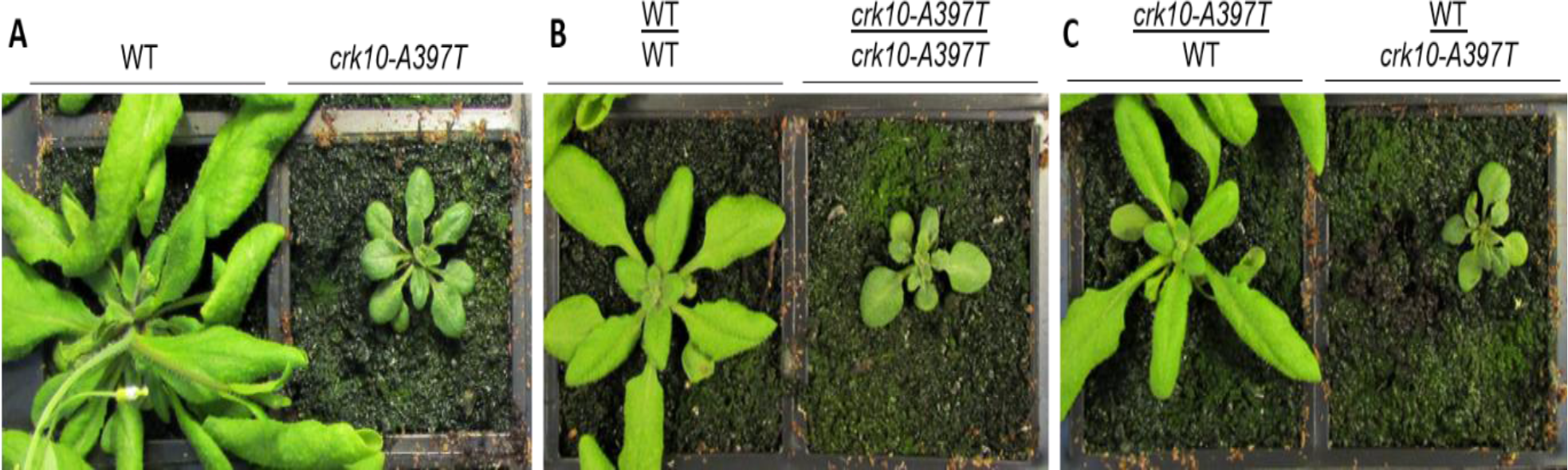
The root-hypocotyl system is responsible for the dwarf phenotype of *crk10-A397T* mutant plants. Images of non-grafted plants (A), self-graft controls (B) and graft combinations (C) of WT and *crk10-A397T* mutant. Plants were imaged 3 weeks after micrografting was performed. The phenotype observed for the reciprocal grafting combinations were consistently observed in two independent repetitions of the experiment. An average number of 10 grafts per combination was recovered each time. Annotation: scion / rootstock.

### The hypocotyl transcriptome of *crk10-A397T* carries the signature of a plant responding to stress

The effect of the *crk10-A397T* mutation on the transcriptome was analysed by RNA-Seq. Total RNA was isolated from 15-, 21- and 35-day-old WT and mutant hypocotyls, which corresponds to the time points used for morphological analysis. Principal component analysis (PCA) showed good clustering of replicates according to genotypes and developmental time points (Supplemental Figure S11A). Following normalization and statistical analysis of the sequencing results (q ≤ 0.05; log_2_ fold change threshold of ±1), we obtained 523 (15 days), 1836 (21 days) and 913 (35 days) differentially expressed genes (DEG), of which 274 were common to all time points. These DEGs were selected as the core set and taken forward for analysis (Figure 6A; Supplemental Figure S11B; Supplemental Tables S1-S4). Comparison to public datasets using the GENEVESTIGATOR® Signature tool (Hruz et al., 2008) revealed that the transcriptome signature of the *crk10-A397T* mutant was most similar to Arabidopsis plants challenged by fungal and bacterial pathogens (S*clerotinia sclerotorum, Plectosphaerella cucumerina* and *Pseudomonas syringae*) and exposed to abiotic stress (treatment with fenclorim and sulfometuron methyl) (Figure 6C). Equally, Gene Ontology (GO) enrichment analysis of the up-regulated genes within the core set (246 genes) with AgriGO v.20 (Tian et al., 2017; False Discovery Rate < 0.05) revealed that terms associated with the biological functions “Defense response” (GO:0006952; p=2.30E-26), “Response to stimulus” (GO:0050896; p=1.40E-24) and “Response to stress” (GO:0006950; p=2.00E-24) are significantlly overrepresented (Figure 6B; Supplemental Tables S5-S6). In accordance with the whole transcriptome data, marker genes indicative for the activation of the salicylic acid (SA)- and jasmonic acid (JA)-regulated defense pathways, such as pathogenesis-related and camalexin biosynthesic genes (Uknes et al., 1992; Thomma et al., 1998; Ahuja et al., 2012), are highly up-regulated in the *crk10-A397T* mutant (Supplemental Table S7). Transcription factors belonging to the WRKY, MYB and NAC-domain containing families are prominent among the regulatory genes induced by *crk10-A397T,* many of which have been associated with the modulation of stress responses (Supplemental Table S8). Therefore, biotic and abiotic stress-responsive pathways are constitutively activated in the *crk10-A397T* mutant, supporting our hypothesis that *crk10-A397T* is a gain-of-function allele of *CRK10*.

**Figure 6.**
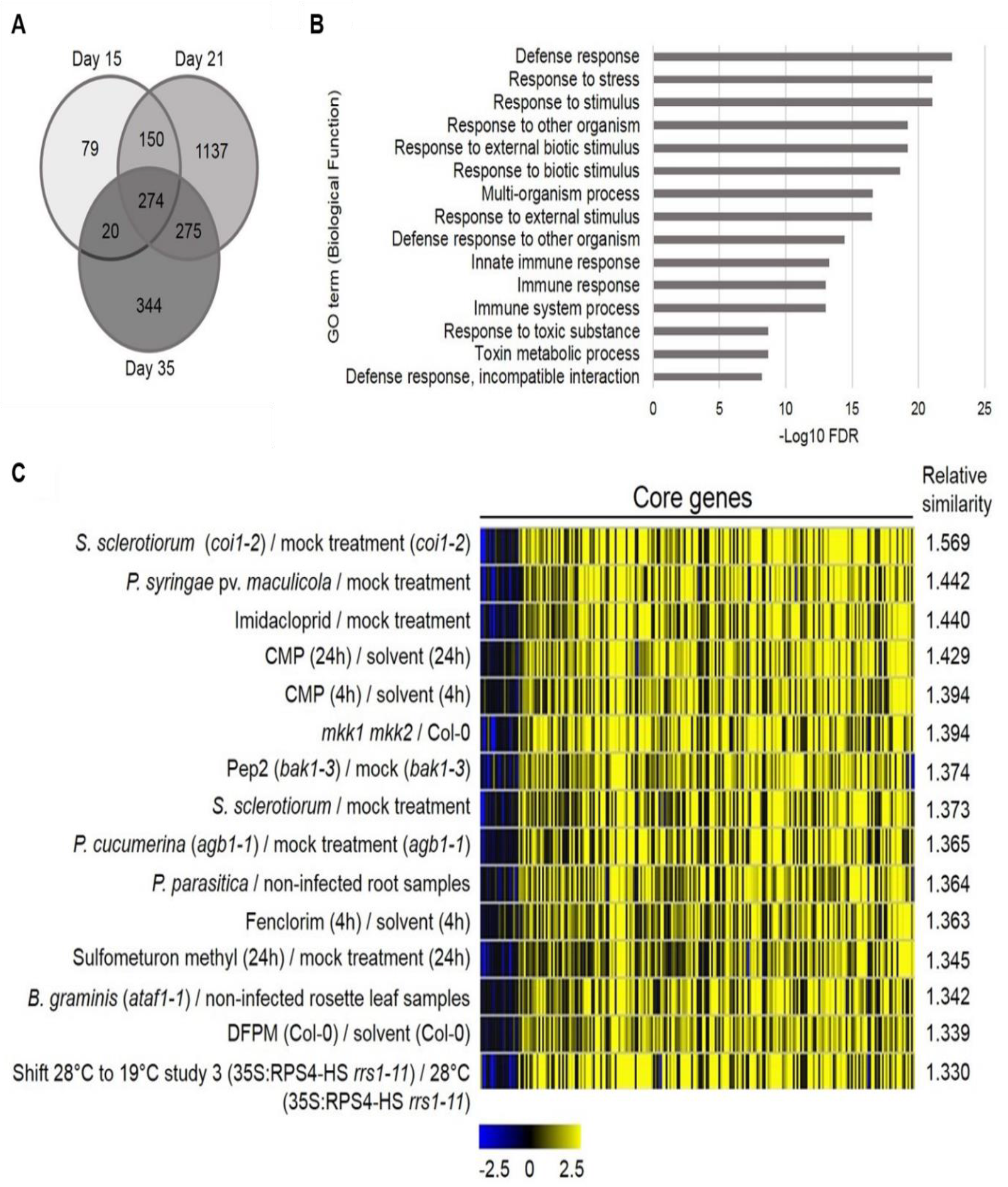
Transcriptional reprogramming in the *crk10-A397T* mutant shows activation of defense responses to biotic and abiotic stresses. A, Venn diagram displaying the number of differentially expressed genes (DEGs) in the *crk10-A397T* hypocotyls compared to the WT at each developmental time point. B, Top 15 enriched GO terms (Biological Process) for the up-regulated core DEGs in the *crk10-A397T* mutant plotted against their respective −log_10_ FDR. C, Top 15 perturbations showing highest overall similarity to *crk10-A397T* mutant expression signature (analysis performed using the GENEVESTIGATOR® SIGNATURE tool; log2 fold change values of time point 21 days was used as input for 274 core genes).

Furthermore, analysis of the lists of DEGs for each individual time point revealed that, specifically at 21 days, genes involved in the biosynthesis, signalling and homeostasis of the hormone abscisic acid (ABA) were also significantly up-regulated in the *crk10-A397T* mutant (Supplemental Table S9).

### *crk10-A397T* mutant hypocotyls contain increased levels of the stress hormones salicylic acid (SA) and abscisic acid (ABA)

In order to corroborate the transcriptomic data we quantified the levels of the stress hormones salicylic acid (SA), abscisic acid (ABA) and jasmonic acid (JA) from the hypocotyls of 3-week-old WT and *crk10-A397T* plants (Supplemental Table S10). In agreement with the transcriptional induction of stress-responsive pathways, the levels of free SA and ABA were increased approximately 3 and 1.5 times, respectively, in the mutant hypocotyls. In contrast, JA levels were not significantly different to the WT.

### The *crk10-A397T* mutant displays enhanced resistant to infection by the vascular pathogen *Fusarium oxysporum* f. sp. *conglutinans* 699

With defense responses constitutively up-regulated in the mutant, we next wanted to investigate whether this is reflected in an enhanced resistance to pathogens. Since *CRK10* expression is detected mainly in the vasculature, we chose the vascular pathogen *Fusarium oxysporum* f. sp. *conglutinans* 699, an isolate known to infect Arabidopsis for the assay (Masachis et al., 2016). *CRK10* OE-1 and *crk10-2* lines were also included in the experiment as overexpression and knockout of other CRKs often showed enhanced/decreased resistance phenotype to pathogens (Acharya et al., 2007, Yeh et al., 2015; Yadeta et al., 2017). The progression of the infection was observed (Figure 7A-D) and a time-mortality curve was recorded from 7 to 20 days post inoculation (Figure 7E). Our results showed that the susceptibility of WT and *crk10-2* plants to the pathogens was very similar, with both genotypes reaching over 65% of mortality at the end of the experiment. *CRK10* OE-1 and *crk10-A397T* plants exhibit a similarly low mortality rate until 10 days post inoculation, although *CRK10* OE-1 plants reach a final death toll of 47.5% in contrast to the lowest overall death count displayed by the *crk10-A397T* mutant of around 18%. Statistical analysis (Deviance test, 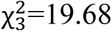, p < 0.001) confirmed the differences in the probability of survival between genotypes, with the *crk10-A397T* mutant having the highest chance of survival of 81.25%, followed by 52.5% for the *CRK10* OE-1 plants and just over 30% for both *crk10-2* and WT (Figure 7F). Fungal burden quantification by qPCR showed increased *F. oxysporum* biomass in WT and *crk10-2* plants compared to *CRK10* OE-1 and *crk10-A397T* mutant at 7 days post inoculation, in agreement with the mortality trend results (Supplemental Figure S12). The experiment was performed twice with similar results. Therefore, our results strongly suggest that *crk10-A397T* mutant plants are more resistant to infection with *F. oxysporum*, reinforcing our hypothesis that the transcriptional responses induced by the *crk10-A397T* mutant allele are effective at reducing the spread of root-infecting vascular pathogens.

**Figure 7.**
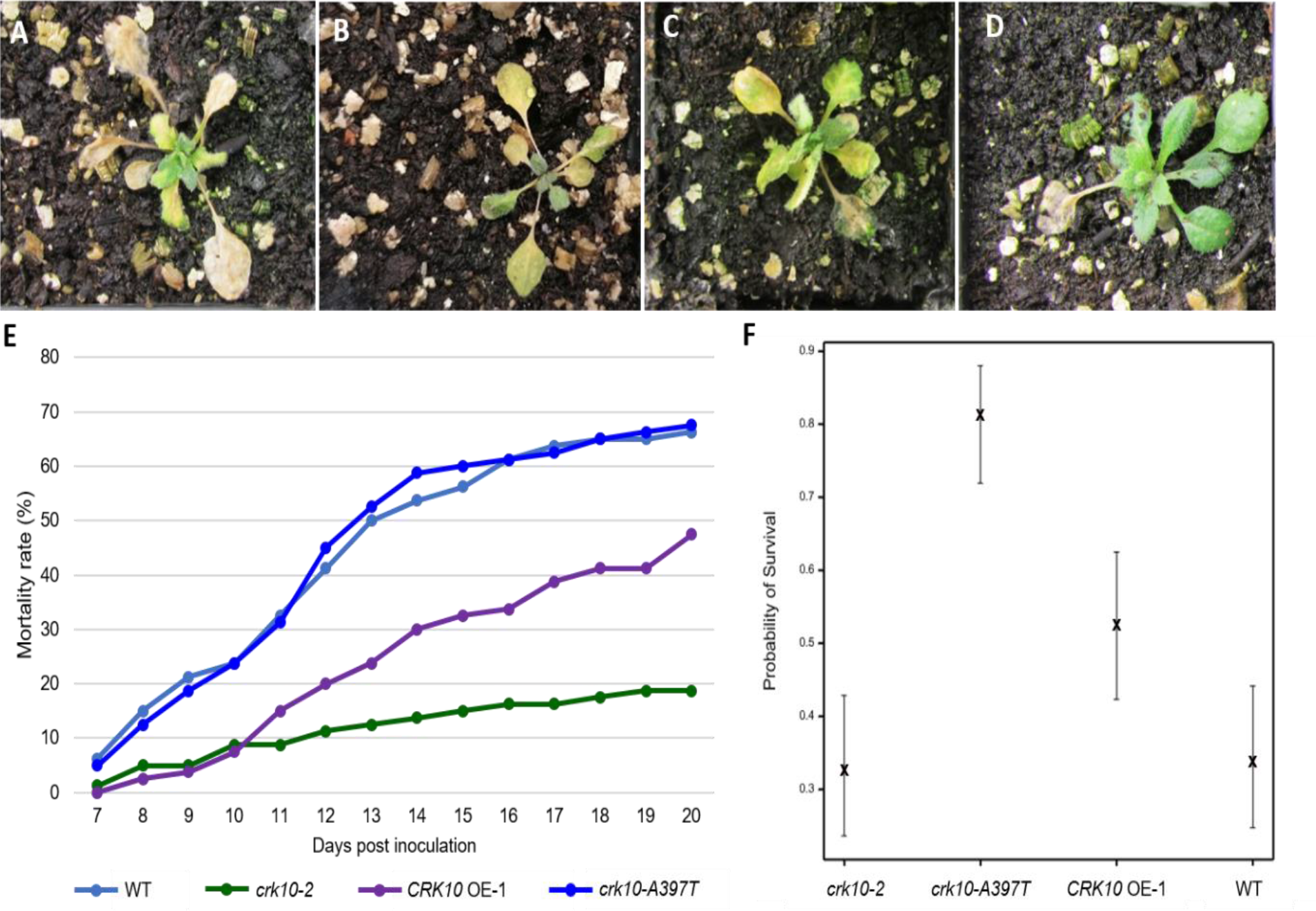
*crk10-A397T* mutant plants are more resistant to infection with *Fusarium oxysporum* f. sp. *conglutinans* 699. A-D, Representative images of WT (A), *crk10-2* (B), *CRK10* OE-1 (C) and *crk10-A397T* mutant (D) plants at 11 days post inoculation with *F. oxysporum*. E, Mortality curve of WT, *crk10-A397T*, *CRK10* OE-1 and *crk10-2* plants from day 7 to 20 post inoculation with *F. oxysporum*. Mortality is shown in percentage. The experiment was repeated twice with similar results. F, Probability of survival of each genotype following inoculation with the pathogen *F. oxysporum*. Associated 95% confidence intervals are shown. Deviance test, 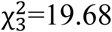, p < 0.001.

## DISCUSSION

Previous efforts to assign functions to specific CRKs mainly focused on the characterisation of T-DNA and overexpression lines. However, being a large multigene family, effects of knocking out one specific member are often masked by redundancy and, in the absence of known stimulants, overexpression lines are generally phenotypically indistinguishable from WT. Here we report the characterisation of the *crk10-A397T* mutant which harbours a gain-of-function allele of CRK10, to our knowledge the first such mutant obtained for this class of receptors in *Arabidopsis*.

### The crk10_A397T mutation lies in an important regulatory subdomain of the kinase domain of CRK10

The point mutation responsible for the conversion of CRK10 into a gain-of-function allele (alanine 397 by threonine) lies in kinase subdomain III, right at the C-terminus of the αC-helix and at the start of the short αC-β4 loop which links the αC-helix to the b strand 4. The importance of this region for kinase regulation has been studied in numerous mammalian kinases with mutations residing in this area often leading to kinase de-regulation and disease, but equivalent studies in plant kinases are absent. The combined αC-helix and αC-β4 loop has been shown to be a critical allosteric docking site which plays a crucial part in kinase regulation (Yeung et al., 2020). As the 3D model of the CRK10 kinase domain suggests, the substituted amino acid lies on that surface of the helix, which faces away from the active site of the kinase and which is known to provide an interface for interactions with many regulatory domains and proteins. For human CDK2, for example, it was shown that residue Lys-56, equivalent to the position of Thr-397 in *crk10-A397T,* is involved in its interaction with cyclin necessary for the stabilisation of the active form of the kinase (Jeffrey et al., 1995), whereas in the case of the Ser/Thr-protein kinase B-Raf (BRaf), where dimerisation is thought to be an important part of the activation mechanism, the αC-Helix/αC-β4 loop provides the interface for dimer formation (Rajakulendran et al., 2009). This region can also function as a *cis*-regulatory site as shown for the human leucine-rich repeat kinase 2 (LRRK2), where it provides a firm docking site for the C-terminal residues of the kinase which keeps it in an inactive conformation (Deniston et al., 2020, Taylor et al., 2020). As the αC-helix/αC-β4 loop is a feature common to all eukaryotic kinases, it seems reasonable to speculate that the amino acid substitution in *crk10-A397T* perturbs the interaction of CRK10 with a regulatory partner. It is also noteworthy that the αC helix and αC-β4 loop is a highly conserved region within the family of CRKs and only three residues are found to occupy the position equivalent to Ala-397 of CRK10 among all the members: alanine, threonine or serine (Supplemental Figures S13 and S14). It now remains to be seen whether other members of the CRK-family are similarly affected by an analogous mutation in their αC-helix/αC-β4 loops.

### Threonine 397 in the kinase domain of CRK10 is an auto-phosphorylation site *in****situ***

*In situ* phosphorylation analysis of the cytoplasmic kinase domains of WT and mutant CRK10 by LC-MS/MS determined that CRK10, being an RD kinase, shows the typical auto-phosphorylation pattern of conserved phosphorylation sites in the activation loop, on which the activity of this class of kinases is dependant (Nolen et al., 2004, Beenstock et al., 2016). Thr-507 and Ser-508 were identified as unambiguous phosphorylation sites, with Ser-508 likely to be functionally equivalent to Thr-450 of BAK1, which plays a key role in its activation by maintaining the active conformation of the activation loop (Yan et al., 2012). However, we also detected phosphorylation of tyrosine residues (Tyr-363 and Tyr-514), with Tyr-514 residing in the activation loop, which establishes CRK10 as a dual specificity kinase. Additional phosphorylated residues were detected in the juxtamembrane domain (Thr-340) and at the C-terminus (Thr-625, Ser-662 and Thr-664). Phosphorylation of residues in these non-catalytic regions have been shown to play an important role in the recognition and/or phosphorylation of downstream substrates, although these are unknown for CRKs (Wang et al., 2005; Oh et al., 2012; Zhou et al., 2020). The phosphorylation pattern of His-CRK100kd^WT^ and His-CRK10kd^A397T^ was identical, with the notable exception of the substituted Thr-397 functioning as an additional phosphorylation site in His-CRK10kd^A397T^. Based on the experimental evidence, it is currently not possible to distinguish whether Thr-397 *per se* or whether its phosphorylated form is the underlying cause for the phenotypic effects observed in the *crk10-A397T* mutant. The significance of the identified *in situ* phosphorylation sites for CRK10 activity *in vivo* also remains to be determined.

### The dwarf phenotype of the *crk10-A397T* mutant is linked to the collapse of xylem vessels in roots and hypocotyls

Phenotypically, the *crk10-A397T* mutant is a dwarf, which we showed to be the consequence of collapsed xylem elements in the roots and hypocotyl. Collapsed xylem vessels are thought to occur due to alterations in the cell wall composition, leading to cell wall weakening and a consequent inability to withstand the negative pressure generated by transpiration, as reported in the *eskimo* and *irx* mutants, for example (Turner & Somerville, 1997; Lefebvre et al., 2011). Although secondary cell wall defects are generally readily observed when analysed by TEM, we could not detected any obvious differences in the vessel cell wall ultrastructure in cross sections of 3-week-old hypocotyls of the *crk10-A397T* mutant and WT. Considering the severity of the vessel collapse in the mutant this was unexpected and future analysis of the composition of the cell wall will be necessary in order to determine the biochemical defects. Intriguingly, although *CRK10* is expressed in vascular-associated tissues in both stem and hypocotyl as shown by reporter lines and qPCR analysis, xylem elements in the stem seem to develop normally in the mutant. Why xylem elements are defective in one organ but not the other can, therefore, not simply be explained by a restricted expression pattern of CRK10 but, considering that RLKs usually reside in large complexes at the plasma membrane, might be due to tissue-specific composition of the “receptorsomes”. However, grafting experiment strongly suggest that it is the defective vasculature in the belowground organs causing the dwarf phenotype, as WT scions grafted onto *crk10-A397T* rootstocks become similarily stunted whereas *crk10-A397T* scions grafted on WT rootstocks develop normally.

### Stress responses are constitutively activated in the hypocotyl of *crk10-A397T*

Collapsed vessel elements are thought to impede proper water transport which is likely to be perceived as drought stress leading to an increase of ABA. This could explain the elevated levels of ABA which we detected in the hypocotyl of *crk10-A397T* and the up- regulation of genes involved in ABA synthesis, perception and response (Supplemental Table S9). A similar increase in ABA levels has been observed in other cell wall mutants such as *esk1* and *irx1-6* whereas *irx3*, *irx5* and *irx9* contained numerous constitutively up-regulated ABA-responsive genes (Chen et al., 2005; Hernández-Blanco et al., 2007; Lefebvre et al., 2011; Faria-Blanc et al., 2018; Xu et al., 2020). The stunting of these mutants has been suggested to be the consequence of the response to drought signalling hormones resulting in the suppression of growth, which could also explain the dwarfism of *crk10-A397T*. It will be interesting to investigate whether dwarfing of *crk10-A397T* can be alleviated by an ABA insensitive mutant such as *abi1* (Koornneef et al., 1984).

The severe collapse of the xylem elements in the roots and hypocotyl of *crk10-A397T* is most likely due to alterations of the composition of the cell walls and it is increasingly being reported that modification of cell wall composition by genetic or chemical means leads to the constitutive activation of defense pathways and an altered resistance to pathogens. The primary cell wall mutant *ixr1/cev1*, for example, with reduced crystalline cellulose content due to the defective cellulose synthase CESA3, displays constitutive activation of jasmonic acid and ethylene signalling (Ellis et al., 2001; Ellis et al., 2002). Transcriptomic data obtained for the secondary cell wall mutants *irx1-6*, *irx5-5* and *irx9* showed that defense-related genes are constitutively expressed in these mutants (Hernández-Blanco et al., 2007; Faria-Blanc et al., 2018). In line with these reports, our data showed that the signature of the core set of DEGs of *crk10-A397T* was most similar to the transcriptome of plants responding to biotic and abiotic stress. Canonical SA-dependent marker genes (PR1, PR2 and PR5) and genes involved in the synthesis of the tryptophan-derived antimicrobial compounds (camalexin, glucosinolates) are significantly up-regulated in the mutant, as are numerous transcription factors usually associated with coordinating stress responses (Supplemental Tables S7 and S8). Concomitant with gene induction, SA levels in the *crk10-A397T* mutant hypocotyls are increased, whereas changes in JA levels were not significant. Induction of defense pathways due to cell wall impairment manifests itself frequently in alteration of disease resistance to a wide range of pathogens (Houston et al., 2016; Bacete et al., 2018). Molina et al. (2021), for example, showed that from a panel of 34 cell wall mutants affecting a wide range of different cell wall compounds, 29 had an altered, mainly enhanced, resistance response to pathogens comprising different parasitic lifestyles. In order to assess disease resistance of *crk10-A397T* we chose the root-infecting, hemi-biotrophic vascular wilt pathogen *F. oxysporum* to perform a pathogen assay, bearing in mind the vasculature associated expression of *CRK10*. In agreement with the studies linking cell wall modification to altered disease resistance, we found that the *crk10-A397T* mutant was significantly more resistant to the pathogen, part of which could be due to the fact that collapsed xylem vessels act as physical barriers slowing pathogen progression in the roots.

## Conclusion

Taken together, the experimental evidence provided in this report suggests that *crk10-A397T* is a mutant which contains a gain-of function allele of the cysteine rich receptor-like kinase CRK10. The transcriptome of the mutant carries the typical signature of a plant responding to stress, although at this stage it is not possible to differentiate whether the stress response is the direct consequence of spontaneous signalling occurring because of the mutated CRK10 receptor or whether it is the response to the perception of the defective xylem vessel cell walls. The mutation lies in the αC-Helix/αC-β4 loop which is emerging as a key structural element in the regulation of kinase function and regulation and a hot-spot for post-translational modifications (Yeung et al., 2020). It is noteworthy that the only other semi-dominant mutant reported for a CRK in rice, *als1*, which develops spontaneous lesions on leaf blades and sheaths also localises to this loop (Du et al., 2019). Future work will now need to address the consequence of the *A397T* mutation for kinase activity. The dwarf and collapsed xylem phenotype of the *crk10-A397T* mutant could, for example, be caused by CRK10 having been converted into a constitutively active kinase and thereby reflect the function of wild type CRK10. However, another scenario which needs to be taken into consideration is that CRK10 A397T has assumed a novel function and interferes with an endogenous pathway distinct from the normal mode of action of wild type CRK10.

## MATERIALS AND METHODS

### Plant Materials and Growth Conditions

*Arabidopsis thaliana* ecotype Col-0 plants were grown in Grobanks cabinets (CLF 2006, Plant Climatics, Germany) in Levington F2 + Sand compost in long day conditions (16h/8h), 23/18 °C day/night temperature, 200 µmol m^-2^ sec^-1^ of light intensity. For *in vitro* experiments, surface sterilized seeds were placed on ½ MS plates (Murashige & Skoog, Duchefa Biochemie).

### RNA-Seq

Total RNA was extracted from hypocotyls of Arabidopsis plants using the RNeasy Mini Kit (QIAGEN), and samples were subsequently treated with DNase Turbo DNA-free kit (Thermo Fisher Scientific). Each biological replicate consisted of a pool of 50-60 hypocotyls, and four biological replicates were isolated per genotype per time point. RNA quality was assessed on a 2100 Bioanalyzer (Agilent Technologies). Library preparation and paired-end sequencing, Illumina HiSeq 125 PE sequencing was performed by Exeter Sequencing Service (University of Exeter, UK).The Rothamsted instance of the Galaxy (https://usegalaxy.org/) bioinformatics web pipeline was used to perform quality control (MultiQC; https://multiqc.info/), trimming (Trimmomatic; Bolger et al., 2014) and mapping to the reference genome (HiSAT2; Kim et al., 2015). The table of counts was acquired using the featureCounts functions in the Subread (Liao et al., 2019; https://bioconductor.org/packages/release/bioc/html/Rsubread.html) on the R Bioconductor platform (https://bioconductor.org/). Genes with less than 3 samples with counts ≥5 were discarded. Differentially expressed genes were identified (using the default Wald test) in the R (v3.6.1) Bioconductor package DESeq2 (Love et al., 2014; https://bioconductor.org/packages/release/bioc/html/DESeq2.html). Gene Ontology (GO) enrichment analysis (Single Enrichment Tool, AgriGO v2.0; Tian et al., 2017) and similarity comparison with deposited micro-array datasets (Signature tool, GENEVESTIGATOR®, Hruz et al., 2008) were performed for the set of 274 core differentially expressed genes.

### Quantitative PCR

RNA was isolated using TRI Reagent® (Merck) and treated with DNase I, Amplification Grade (Thermo Fisher Scientific) prior to cDNA synthesis with SuperScript™ III Reverse Transcriptase (Thermo Fisher Scientific). All qPCR reactions were performed in a LightCycler® 96 Real-Time PCR System (Roche Diagnostics) using the FastStart Essential DNA Green Master (Roche Diagnostics). *ACTIN2* (AT3G18780) and *UBC21* (AT5G25760) were used as internal controls (Supplemental Table S11), and the 2^−ΔΔCt^ method (Livak & Schimittgen, 2001) was used to calculate relative expression.

### Quantification of plant hormones

Hypocotyl samples (75 – 100 mg fresh weight) isolated from 3-week-old plants were used for the quantification of hormones as described in Camut et al., 2019. A 2.6 µm Accucore RP-MS column (100 mm length x 2.1 mm i.d.; ThermoFisher Scientific) and a Q-Exactive mass spectrometer (Orbitrap detector; ThermoFisher Scientific) were used. The concentrations of hormones in the extracts were determined using embedded calibration curves and the Xcalibur 4.0 and TraceFinder 4.1 SP1 programs. Three biological replicates per genotype were analysed.

### Cloning of genetic constructs

The list of primers used in this study can be found in Supplemental Table S11. To generate the construct for the complementation of the *crk10-A397T* mutant and overexpression of *CRK10* in the WT plants, the cDNA sequence of *CRK10* was amplified from the cDNA clone U60398 with *Sal*I and *Sac*I restriction sites and cloned into pJD330 downstream of the 35S promoter. The construct 35S:*CRK10*-NOS was then amplified with *Asc*I restriction sites and cloned into the RS 3GSeedDSRed MCS vector. To obtain the *CRK10*_Pro_:*crk10-A397T*-NOS construct, the 1 kb genomic sequence upstream of *CRK10* (*CRK10*_Pro_) was amplified from Arabidopsis Col-0 genomic DNA with *Sph*I and *Sal*I restriction sites, and was cloned in pJD330 upstream of the *CRK10* cDNA sequence. The G>A mutation harboured by the *crk10-A397T* mutant was introduced by *in vitro* mutagenesis.

The *CRK10*_Pro_ sequence was amplified with *Sph*I and *Nco*I restriction sites, and the fragment was ligated upstream of the *β-glucuronidase* (*GUS*) gene in pJD330. The *CRK10*_Pro_:*GUS*-NOS construct was amplified with *Asc*I restriction sites and cloned into the RS 3GSeedDSRed MCS vector. To obtain a translational fusion of CRK10 with the fluorescent protein mCherry, the stop codon of *CRK10* was removed and a *Sac*I restriction site was introduced by *in vitro* mutagenesis between *CRK10* and the NOS terminator sequence in pJD330. The coding sequence of *mCherry* was cloned with *Sac*I restriction sites on both ends and ligated in frame with *CRK10*. The construct *CRK10-mCherry*-NOS was then amplified with *Sal*I and *Not*I restriction sites and cloned into pENTR1A Dual Selection Vector (Invitrogen). Following LR reaction (Gateway) with the binary vector pB2GW7 (Invitrogen), the construct 35S:*CRK10-mCherry*-NOS was obtained. Finally, to generate the constructs for recombinant protein in *E. coli*, the coding sequence of the kinase domain of *CRK10* was amplified with *Sal*I and *Not*I restriction sites and cloned into pENTR1A Dual Selection Vector (Invitrogen) prior to recombination with pDEST17 (Invitrogen), where it was cloned in frame with an N-terminal 6x His-tag. I*n vitro* mutagenesis was used to generate the gain-of-function mutation of CRK10 (A397T) as well as the dead kinase variant by mutation of the invariant aspartic acid in the active site to an asparagine (D473N). All constructs used for the generation of transgenic plants were introduced in A. tumefaciens cells strain GV3101 using the freeze and thaw protocol (An et al., 1988), and the floral dip method (Clough and Bent, 1998) was used to transform Arabidopsis plants.

### Transient expression of fluorescent fusion in *N. benthamiana*

Protocol was adapted from Sparkes et al., 2006. Transient expression of the fusion protein was observed 72 hours post-infiltration.

### Microscopy

For the preparation of thin sections for light microscopy, plant tissue samples were fixed in 4% paraformaldehyde / 2.5% glutaraldehyde (0.05 M phosphate buffer pH 7.2), dehydrated in graded ethanol series (10 − 100%) and infiltrated with LRWhite resin (Agar Scientific). Sections were prepared using a Reichert Ultramicrotome (section thickness: 1-2 µm) and stained with 0.5% potassium permanganate. Thin sections were observed with a Zeiss Axiophot equipped with a Q-Imaging Retiga EXi CCD mono digital camera with RGB filter wheel (QImaging, Canada) coupled with the Metamorph software (Molecular Devices, USA). Non-stained resin embedded thin sections were also used for the detection of the auto-fluorescence of lignin using the confocal microscope (excitation: 405 nm; emission: 451-480 nm and 560-612 nm). For the detection of mCherry fluorescence from *N. benthamiana* leaves and hypocotyls of Arabidopsis, a small piece of fresh tissue sample was mounted on a glass slide with glass cover slip and water, and the fluorescence was detected using the confocal microscope (excitation: 561 nm laser; emission: 578 - 639 nm). Plasmolysis was performed using a 0.8M mannitol solution for 40 minutes and images acquired prior to and after treatment. Fluorescent signals were detected from thin sections and whole plant samples using a Zeiss 780 LSM confocal microscope. For the preparation of samples for transmission electron microscopy (TEM), plant material was fixed by high-pressure freezing in 0.1 M sucrose using a Leica HPM100, followed by freeze substitution with 100% ethanol (Leica EM Auto Freeze substitution) and infiltration with LRWhite resin (Agar Scientific). Ultra-thin sections prepared with a Leica EM UCT Ultramicrotome (section thickness: 90 nm) were collected on pioloform/carbon-coated nickel grids (Agar Scientific) and stained with 2.5% uranyl acetate and Reynolds lead citrate (Reynolds, 1963). Ultrathin sections were imaged using a JEOL-2100Plus Transmission Electron Microscope (JEOL, Japan) equipped with a Gatan OneView IS Camera (Gatan, USA).

### Micrografting of Arabidopsis seedlings

Grafting protocol was performed as previously described (Turnbull et al., 2002).

Successful grafts were transferred to soil 7-10 days post-grafting.

### GUS staining

Plant tissue was incubated overnight in X-Gluc (Melford) solution at 37 °C. Chlorophyll was removed with 80% ethanol prior to imaging using a Leica M205 FA Stereomicroscope (Leica Microsystems).

### Recombinant protein expression, purification and analysis

BL21 AI One-Shot E. coli cells (Invitrogen) were used for the expression of recombinant 6x His-tag CRK10 kinase domain variants. Singles colonies were inoculated in liquid media supplemented with antibiotics and 0.1% glucose and grown overnight at 37 °C shaking at 225 rpm. A 250 μL aliquot of the overnight culture was used to inoculate 50 mL of fresh selective media, and cells were grown until OD_600_ = 0.4-0.5 was obtained. L-arabinose was added to the cultures to a final concentration of 0.2% to induce expression of the recombinant protein, and cultures were grown for an additional 3 hours. Cells were harvested by centrifugation (5,000 g x g for 10 minutes), resuspended in equilibration buffer (50 mM sodium phosphate pH 8.0, 0.3 M sodium chloride) and lysed by sonication. The cell lysate was centrifuged for 30 minutes at 5,000 x g and the supernatant was recovered and mixed with HIS-Select Nickel Affinity Gel (Sigma Aldrich) for protein purification (manufacturer’s instructions were followed). The concentration of the purified protein extracts was measured using the Bradford method (Protein Assay Dye Reagent, Bio-Rad). Purified His-tagged proteins were resolved by SDS-PAGE (NuPAGE 4-12% Bis-Tris Protein Gels, Invitrogen) and gels were stained with Quick Coomassie Stain (Generon). Alternatively, proteins were transferred to a PVDF membrane (iBlot Transfer Stack, PVDF, Invitrogen) for Western blotting. The membrane was blocked with 5% powder milk in PBS-T, and incubated with His-probe (H-3) HRP monoclonal antibody (Santa Cruz Biotechnology) for 1 hour. The membrane was washed and incubated with Amersham ECL Western Blotting Detection Reagent (GE Healthcare) and developed in the dark. Protein extracts were also treated with Lambda Protein Phosphatase (Lambda PP, New England BioLabs) according to the manufacturer’s protocol. Briefly, 5 μg of purified protein were incubated with Lambda PP for 1h30min at 30 °C. Phosphatase-treated samples were then resolved on SDS-PAGE and detected by Western blotting.

### Liquid chromatography – Mass Spectrometry/Mass Spectrometry (LC-MS/MS) and MASCOT database search

Gel bands were transferred into a 96-well PCR plate. Peptide bands were cut into 1mm2 pieces, destained, reduced (DTT) and alkylated (iodoacetamide) and subjected to enzymatic digestion with sequencing grade trypsin (Promega, Madison, WI, USA) overnight at 37°C. After digestion, the supernatant was pipetted into a sample vial and loaded onto an autosampler for automated LC-MS/MS analysis. All LC-MS/MS experiments were performed using a Dionex Ultimate 3000 RSLC nanoUPLC (Thermo Fisher Scientific Inc, Waltham, MA, USA) system and a QExactive Orbitrap mass spectrometer (Thermo Fisher Scientific Inc, Waltham, MA, USA). Separation of peptides was performed by reverse-phase chromatography at a flow rate of 300 nL/min and a Thermo Scientific reverse-phase nano Easy-spray column (Thermo Scientific PepMap C18, 2 µm particle size, 100A pore size, 75 µm i.d. x 50cm length). Peptides were loaded onto a pre-column (Thermo Scientific PepMap 100 C18, 5 µm particle size, 100A pore size, 300 µm i.d. x 5mm length) from the Ultimate 3000 autosampler with 0.1% formic acid for 3 minutes at a flow rate of 10 µL/min. After this period, the column valve was switched to allow elution of peptides from the pre-column onto the analytical column. Solvent A was water + 0.1% formic acid and solvent B was 80% acetonitrile, 20% water + 0.1% formic acid. The linear gradient employed was 2-40% B in 30 minutes. Further wash and equilibration steps gave a total run time of 60 minutes. The LC eluant was sprayed into the mass spectrometer by means of an Easy-Spray source (Thermo Fisher Scientific Inc.). All m/z values of eluting ions were measured in an Orbitrap mass analyzer, set at a resolution of 70000 and was scanned between m/z 380-1500. Data dependent scans (Top 20) were employed to automatically isolate and generate fragment ions by higher energy collisional dissociation (HCD, NCE:25%) in the HCD collision cell and measurement of the resulting fragment ions was performed in the Orbitrap analyser, set at a resolution of 17500. Singly charged ions and ions with unassigned charge states were excluded from being selected for MS/MS and a dynamic exclusion window of 20 seconds was employed.

### Protein modelling

A structural model of the kinase domain of CRK10 was generated by homology modelling using PyMOD 3.0 with default parameters (Janson et al., 2017). The kinase domain of BRI1 (PDB code 5LPV) (Bojar et al., 2014) was used as template retaining the ATP analogue (phosphoaminophosphonic acid-adenylate ester) but no other heteroatoms.

### Plant infection assay

Root infection assay was modified from Masachis et al., 2016. Arabidopsis seedlings (12-days-old) grown *in vitro* were inoculated by immersing their roots for 20 minutes in a suspension of 1 x 10^6^ microconidia ml^-1^ of *F. oxysporum* f. sp*. conglutinans* 699. Seedlings were transferred to Levington F2 + Sand compost with vermiculite (3:1) and incubated in growth chamber under long day conditions (16/8h), 28/25 °C day/night temperature. Eighty plants per genotype were planted in a randomised blocked tray design, and mortality was assessed daily between 7 and 20 days post-inoculation. For determination of fungal burden, three independent replications of the experiment were performed. Seedlings were inoculated as described above and collected at 2 and 7 days post-inoculation. Each repetition of the experiment included one pool of eight seedlings per genotype which were processed as one biological replicate. Total DNA extraction was performed according to Yu et al., 2019 and the relative amount of fungal DNA was quantified by qPCR using the *F. oxysporum ACTIN1* normalized to the Arabidopsis *ACTIN2* gene. Results were expressed relative to WT at 2 days post inoculation.

### Statistical analysis

Statistical tests were performed using the Genstat software (Genstat for Windows 21st Edition; VSN International, Hemel Hempstead, UK). Student’s t-test were used to assess statistical differences between two variants. To assess whether the pattern of segregation of the dwarf phenotype followed the expected 1:2:1 ratio, the 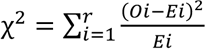 chi-square statistic was used, where O_i_ is the observed count for group i and E*i* is the expected count for group i. Under the null hypothesis of 1:2:1 segregation this test statistic should follow a chi-square distribution with 2 degrees of freedom. The probability of survival of each genotype in the bioassay with *F. oxysporum* was assessed with a generalized linear model (Bernoulli distribution; logit link function fitted to the final mortality outcome of each plant); statistical significance of the genotypic effect was tested after removing variation associated with plant position within rows of different trays and quantified through a Chi-squared statistic of the difference in deviance. Statistical significance of the differences in fungal burden between genotypes were tested by ANOVA (expression levels were log-transformed to meet the ANOVA requirements, and each individual experiment was considered as a block).

### Accession numbers

Gene sequence data for this article can be found in The Arabidopsis Information Resource (TAIR) under the accession number AT4G23180 (*CRK10*). The *CRK10* cDNA clone (U60398) was obtained from the Arabidopsis Biological Resource Center (ABRC). The T-DNA lines SAIL_427_E09 and SALK_116653 were obtained from the Nottingham Arabidopsis Stock Centre (NASC; plants were genotyped according to instructions provided in http://signal.salk.edu). The mass spectrometry proteomics data have been deposited to the ProteomeXchange Consortium via the PRIDE (Perez-Riverol et al., 2019) partner repository with the dataset identifier PXD023831.

## ACKNOWLEDGMENTS

We thank Professor Colin Turnbull (Imperial College London) for help with the micrografting technique; Hannah Walpole, Kirstie Halsey and Dr Eudri Venter (Bioimaging Department, Rothamsted Research) for assistance with light, confocal and transmission electron microscopy; Dr Jason Rudd (Department of Biointeractions and Crop Protection, Rothamsted Research) for invaluable discussions regarding the analysis of the kinase domain of CRK10; Dr Kim Hammond-Kosack (Department of Biointeractions and Crop Protection, Rothamsted Research) for insightful discussions regarding CRK10’s role in plant-pathogen interactions; Kirsty Hassall (Department of Computational and Analytical Sciences, Rothamsted Research) for support with experimental design and statistical analyses; and Professor Antonio di Pietro for providing the fungal isolate used in this study. We thank the University of Exeter for the Exeter International Excellence Scholarship which supported MP while this work was developed. This work was supported by UK Biotechnology and Biological Sciences Research Council grants BBS/E/C/000I0420 and BB/P016855/1. This research was done under the regulation described in the DEFRA license number 101948/198285/6.

